# Network-based hierarchical population structure analysis for large genomic datasets

**DOI:** 10.1101/518696

**Authors:** Gili Greenbaum, Amir Rubin, Alan R. Templeton, Noah A. Rosenberg

## Abstract

Analysis of population structure in natural populations using genetic data is a common practice in ecological and evolutionary studies. With large genomic datasets of populations now appearing more frequently across the taxonomic spectrum, it is becoming increasingly possible to reveal many hierarchical levels of structure, including fine-scale genetic clusters. To analyze these datasets, methods need to be appropriately suited to the challenges of extracting multilevel structure from whole-genome data. Here, we present a network-based approach for constructing population structure representations from genetic data. The use of community detection algorithms from network theory generates a natural hierarchical perspective on the representation that the method produces. The method is computationally efficient, and it requires relatively few assumptions regarding the biological processes that underlie the data. We demonstrate the approach by analyzing population structure in the model plant species *Arabidopsis thaliana* and in human populations. These examples illustrates how network-based approaches for population structure analysis are well-suited to extracting valuable ecological and evolutionary information in the era of large genomic datasets.

## 1 Introduction

Population structure is ubiquitous in natural populations, and the ability to detect it from information on genetic variation has been instrumental to our understanding of evolution and ecology. By studying population structure, researchers can investigate processes of gene flow and migration, quantify the evolutionary impacts of fragmentation and movement barriers, inform decisions concerning the conservation and management of populations, and interpret selection signals found in genomic studies. Currently, population structure analysis is an essential element in the toolkit of evolutionary biologists, conservation biologists, and ecologists, both for exploratory data analysis and for hypothesis testing (Guillot et al. 2009; Allendorf et al. 2010; Novembre and Peter 2016).

As genome sequencing technology advances, large-scale whole-genome sequencing is becoming possible not only for describing genetic variation among species, but also for reporting similarities and differences among individuals within populations. Datasets describing genome-wide sequence variation for multiple conspecific individuals are emerging (e.g.,Li et al. 2008; The 1000 Genomes Project Consortium 2015; Alonso-Blanco et al. 2016; Martin et al. 2018; Armstrong et al. 2019), and a surge of whole-genome population datasets is expected in the coming years. Whole-genome population structure analysis presents the promise of high-resolution population structure inference, such as multiple hierarchical levels and detection of fine-scale structure. However, the era of large genomic datasets also introduces challenges for extracting and interpreting population structure at a genomic scale, as well as significant computational difficulties.

Methods such as F-statistics (Excoffier et al. 1992; Holsinger and Weir 2009), principal components analysis (PCA) and other representations of datasets using techniques of multivariate analysis (Menozzi et al. 1978; Jombart et al. 2009), tree-based inference (Bowcock et al. 1994; Pickrell and Pritchard 2012), and STRUCTURE-like methods (Pritchard et al. 2000; Alexander et al. 2009), have been exceedingly successful in enabling inference of population structure from genetic data. Each of these types of methods has distinctive features and disadvantages. F-statistics (e.g. *F*_*ST*_) are easily-communicated summary statistics for measuring genetic differentiation in hierarchically defined population structures, but they require putative groupings of individuals prior to data analysis. Multivariate analysis methods such as PCA do not require a prior grouping of individuals into populations, but they do require visual interpretation or post-analysis clustering in order to place individuals into clusters, and they do not naturally represent hierarchical structure. Tree-based hierarchical analyses depict evolutionary relations between individuals or putatively defined groups (e.g., neighbor-joining, TreeMix; Bowcock et al. 1994; Pickrell and Pritchard 2012); a limitation is that rigid tree-based representations might poorly depict the data when evolution has been particularly non-tree-like. STRUCTURE-like methods cluster individuals into groups based on a pre-defined model inherent to the analysis. Like multivariate analyses, they have the feature that they do not require prior assignment of individuals into groups, and the limitation that a hierarchical representation is not intrinsic to the approach. Multiple hierarchical levels can be explored in STRUCTURE-like methods by repeatedly clustering individuals using models with different numbers of assumed clusters or subsets of the data. Although hierarchical structure is often observed, identification of multiple hierarchical levels depends on the nature of the structure in the data, and is not integral to the method. We are unaware of any population structure inference method that clusters individuals into groups without placing individuals into populations prior to the analysis, and that as a feature intrinsic to the method detects and represents multiple hierarchical levels in a flexible form.

In recent years, network approaches have been introduced to model and analyze population structure (Dyer and Nason 2004; Rozenfeld et al. 2008; Greenbaum et al. 2016; Greenbaum and Fefferman 2017; Han et al. 2017; Kuismin et al. 2017). In network-based population structure inference, the genetic relations between individuals within the populations are formulated as a network, incorporating the many complexities that result from evolutionary processes in natural populations, with few prior assumptions regarding the biological processes involved in generating these relations. The resulting mathematical construct, the network, is then subjected to analyses developed in network science (Newman 2010) to identify topological characteristics that can be interpreted as population structure. Network approaches have shown promise in the detail with which population structure can be described, as well as in computational efficiency (Greenbaum et al. 2016).

Here, we describe a network-based population structure inference approach that allows for efficient clustering of individuals based on genetic information, and that produces hierarchical population structure diagrams. Cluster representations containing many clusters at different levels in a hierarchy can then assist in evaluation and interpretation of the significance of the population structure detected. We provide examples of analyses of population structure in two large well-studied genomic datasets, in *Arabidopsis thaliana* (Durvasula et al. 2017) and in humans (Li et al. 2008), emphasizing the applicability of our approach for studying ecology and evolution.

## 2 Results

### 2.1 Detecting hierarchical population structure using networks

The problem of detecting population structure from genetic data has a particular emphasis on the hierarchical nature of many natural population structures. In order to use networks to detect population structure, we first formulate genotype data as a genetic similarity network, with individuals as nodes and inter-individual genetic similarities as edges. To construct a genetic similarity network, we use a frequency-weighted allele-sharing similarity measure (Greenbaum et al. 2016), with the weighting scheme designed to increase the contribution assigned to shared rare alleles. Constructing pairwise genetic similarity networks for large numbers of loci is a computationally intensive process; however, it is also highly parallelizable and can therefore be completed in reasonable time with the readily-available computation clusters available today, even for whole-genome datasets (see *Computational efficiency* section in *Methods*).

In our approach, the problem of detecting population structure — groups of individuals sharing common evolutionary histories — is equated with the problem of detecting dense subnetworks within the genetic similarity network. In other words, we identify groups of individuals that are highly interconnected. In network science, such dense groups are termed “communities,” and many community-detection methods have been developed in recent years (Newman 2002; Newman 2006; Fortunato 2010).

The genetic similarity network summarizes genetic variation from across the entire genome. However, the network is expected to be extremely dense, with most pairs connected, because any two individuals, even if belonging to distinct subpopulations, are expected to share some alleles inherited from distant common ancestors. This property adds much noise to any signal of structure, potentially decreasing the ability to detect fine-scale structure. In particular, fine-scale population structure is characterized by smaller groups of individuals that share a more recent genetic history within the group than do larger groups representing coarser-scale structure. Therefore, genetic similarity within fine-scale clusters, which are expected to contain only strong genetic similarities, is expected to be higher than that in coarser clusters, which are expected to contain weaker but nontrivial network connections. Consequently, we explore population structure across many hierarchical scales by iteratively pruning weak edges from the network and applying community detection procedures (Fig. 1). By removing weak edges from a coarse cluster, the population structure signals characteristic of fine-scale clusters within the coarser cluster become distinguishable, and another hierarchical level can be revealed. The resulting set of clusters can be described as a tree-like hierarchy of nested clusters, which we term a “Population Structure Tree” (PST). In a PST, the root cluster contains all individuals, the leaf clusters describe the finest-scale structure, and the clusters represented by internal nodes describe intermediate levels of clustering (Fig. 1). The internal clusters overlap with clusters emerging from them as “descendants,” while all the leaf clusters are non-overlapping. Note that in PSTs, no assumptions are made regarding the evolutionary processes that give rise to the hierarchical structure, and the branches need not represent groups that have evolved in isolation.

**Figure 1:**
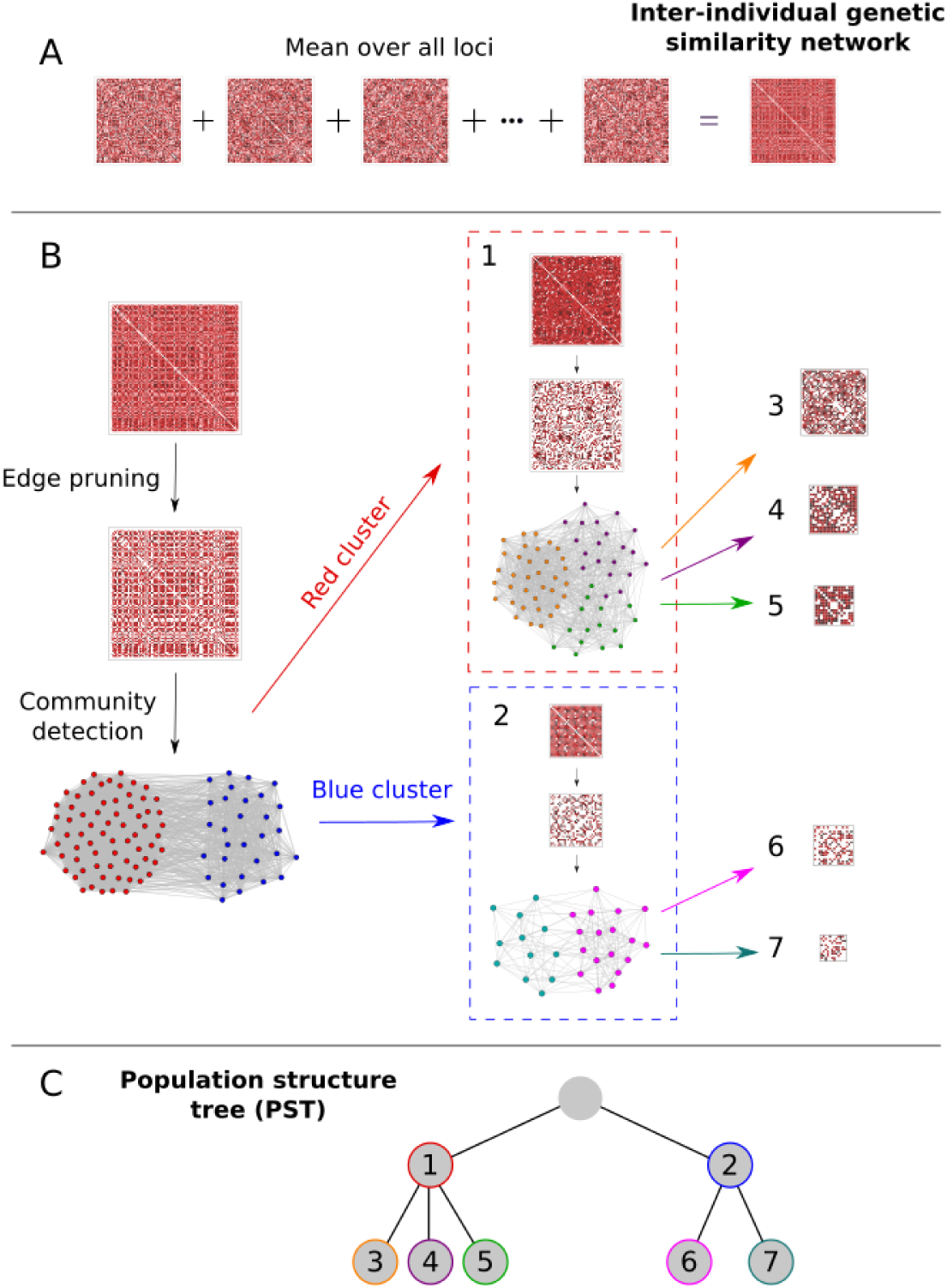
Schematic representation of a network-based construction of a population structure tree (PST) from genomic data. (A) For each SNP, an inter-individual genetic similarity network (adjacency matrix) is constructed using a frequency-weighted allele-sharing genetic similarity measure (Eq. 1). To produce a genome-wide genetic similarity matrix, the mean over all loci is taken. (B) Weak edges are pruned from the matrix, by setting low matrix entries to 0 until a community structure emerges, as detected using network community-detection algorithms. Each community (numbered submatrices) is then analyzed independently in a similar manner. Notice that finer-scale clusters are characterized by darker matrices, indicating structures characterized by higher genetic similarities. (C) The analysis is summarized as a PST diagram, summarizing the hierarchical levels of population structure and their relationships.

### 2.2 Visualization

An important aspect of interpreting population structure analyses is visualization of the outputs. We visualize population structure on geographic maps using a coloring scheme designed to reflect the topology of the inferred PST. We start with a color interval (e.g. “rainbow colors”), and assign it to one of the clusters, which becomes the root cluster for that coloring. Our coloring scheme seeks to describe the topology of the branch connecting the selected root cluster and the leaves. For the selected root cluster, each daughter cluster — a cluster emerging from a parent cluster through the edge-pruning and community detection process — inherits an equal fraction of the color interval, in no particular order. This process of transmission of color intervals to daughter clusters is then repeated until the leaves of the hierarchy are reached on all branches. Each cluster is then assigned a color that corresponds to the midpoint of its assigned color interval. The scheme results in a coloring of a branch of the PST, such that closer clusters in the hierarchy, within and between hierarchical levels, are assigned closer colors on the initial color interval (see examples in Figs. 2–5).

**Figure 2:**
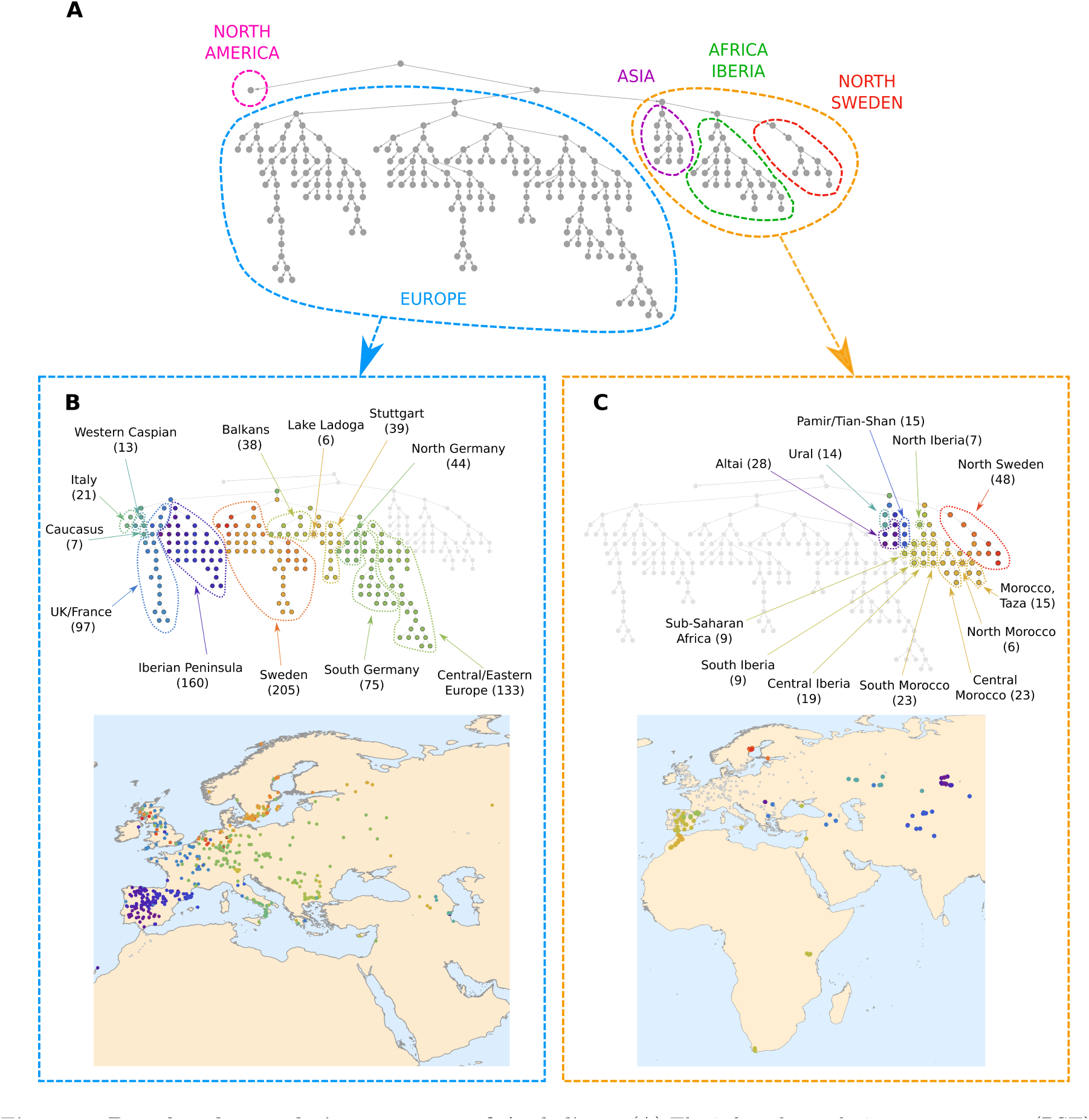
Broad-scale population structure of *A. thaliana*. (A) The inferred population structure tree (PST). Each element in the hierarchy represents a cluster of individuals, and each cluster contains those clusters below it in the hierarchy. The root element represents the entire sample of 1,214 individuals. In colored dashed lines, the main regions corresponding to sampling locations are indicated (with the labels defined *post hoc*). (B) Visualization of the branch corresponding to most European sampling locations. Each cluster is assigned a color such that “closer” colors represent closer clusters in the PST. On the map, each individual is placed at its sampling location and colored according to the finest-scale cluster to which it was assigned. (C) Visualization of the branch corresponding to Africa, Asia, North Sweden, and some samples in the Iberian Peninsula.

**Figure 3:**
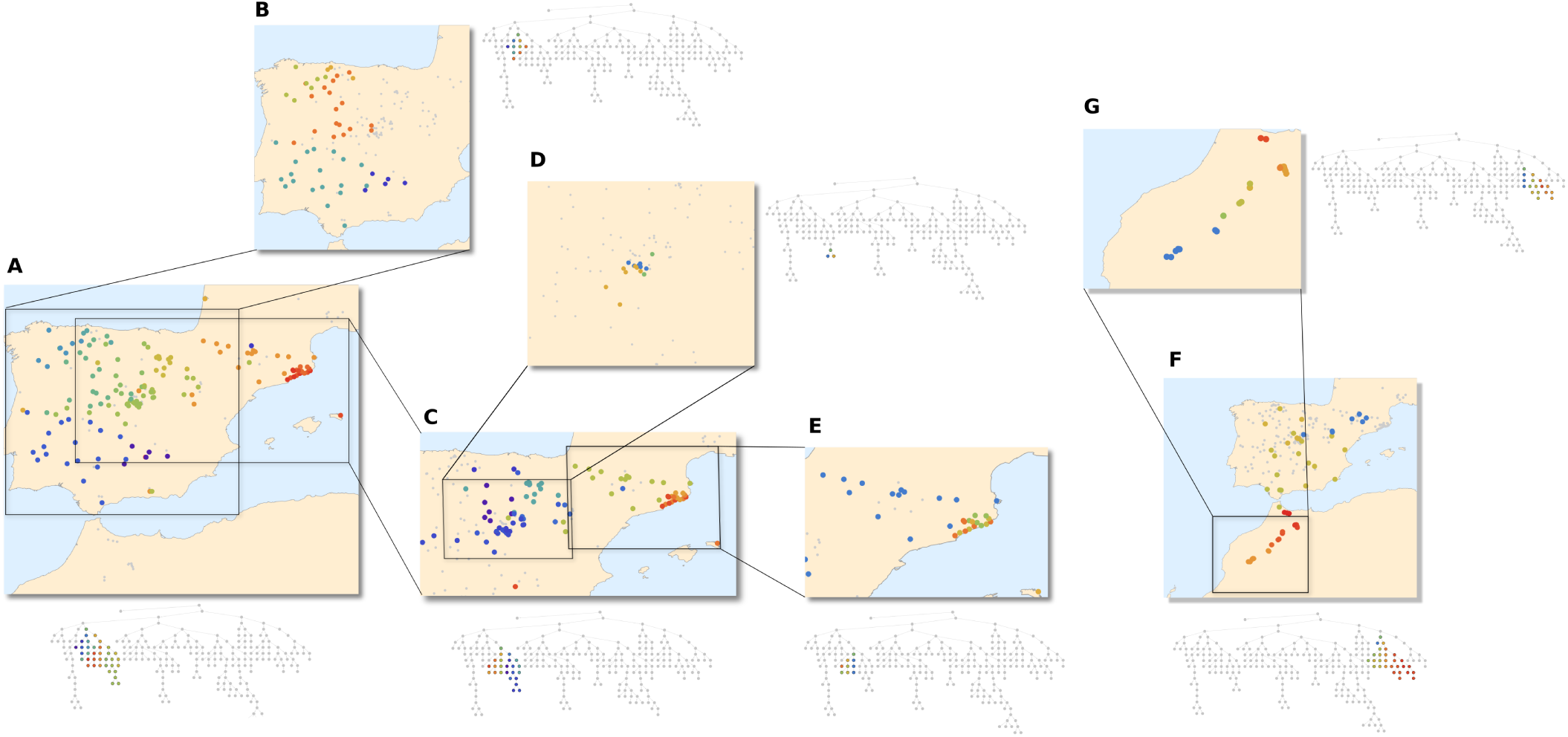
Fine-scale population structure of *A. thaliana* in the Iberian Peninsula and Morocco. On each map, a branch of the inferred PST from Figure 2 is visualized. Adjacent to each map is the PST colored in the same manner as in the map. Panels (A)–(E) show a sub-branch of the primary European branch (blue branch in Fig. 2A), and panels (F) and (G) show a sub-branch of the primarily non-European branch (orange branch in Fig. 2A). (A) Branch corresponding to most of the Iberian population (Iberian peninsula branch in Fig. 2B). (B) Branch corresponding to the western part of the Iberian peninsula, with differentiation along a north-south gradient. (C) Branch corresponding to the northeastern part of the peninsula, with differentiation along an east-west gradient. (D) Branch corresponding to a small area in the center of the peninsula. (E) Branch corresponding to North Spain, with a differentiated sub-branch along the northeast coast. (F) Branch not belonging to the primary European branch in Figure 2A, corresponding to the Iberian Peninsula and Morocco. Population structure extends on a north-south axis from Africa to Europe over the Strait of Gibraltar. (G) Branch corresponding to Morocco, showing fine-scale population structure in Morocco.

**Figure 4:**
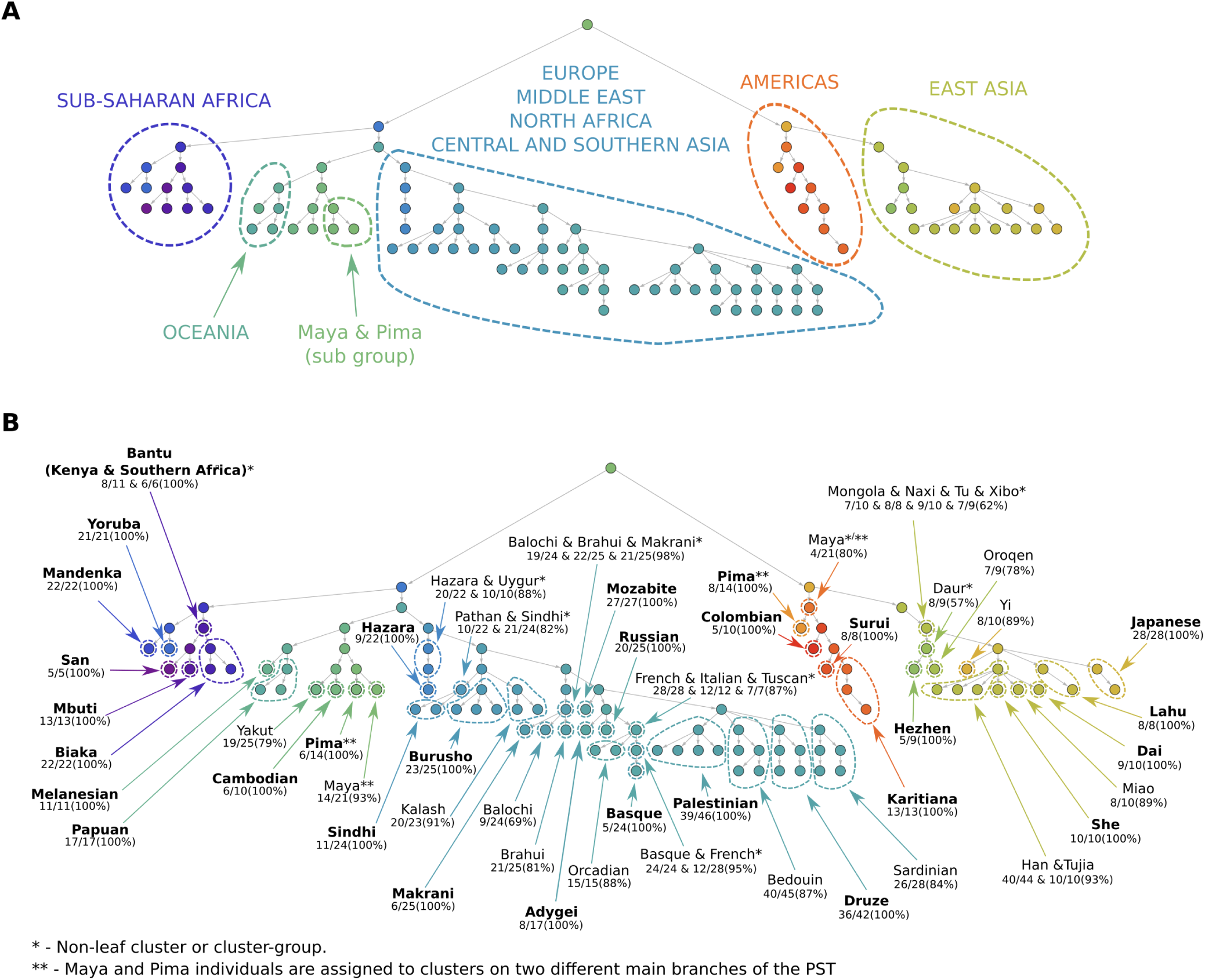
Population structure tree in humans. Closer colors represent closer clusters on the PST. (A) Main branches corresponding to continental groups have been labeled based on assignment of individuals in the branches to population groups (labels defined *post hoc*). Maya and Pima individuals are found in two subgroups, one within the Americas branch and one outside it. (B) Fine-scale structure revealed by the PST. Clusters or cluster-groups in which a majority (> 50%) of individuals have a particular label or labels are circled and marked with the corresponding group labels. Under each label, detailed assignments are given in the format x/y(z%): *x*, number of individuals with the marked label assigned to the cluster-group; *y* number of individuals in the dataset with that label; *z*, proportion of individuals with the marked labels among all individuals assigned to the cluster-group. In the case of cluster-groups with more than one hierarchical level, the detailed assignments refer to the cluster at the highest hierarchical level in the cluster-group. For non-leaf cluster-groups (marked with *), the proportion *z* is taken among all individuals in the cluster-group, omitting all individuals in all descendant clusters assigned a label different from the label of the cluster-group. Labels in bold indicate clusters or cluster-groups (not considering omitted individuals, if relevant) that contain only individuals that have the marked labels (i.e., *z* = 100%).

**Figure 5:**
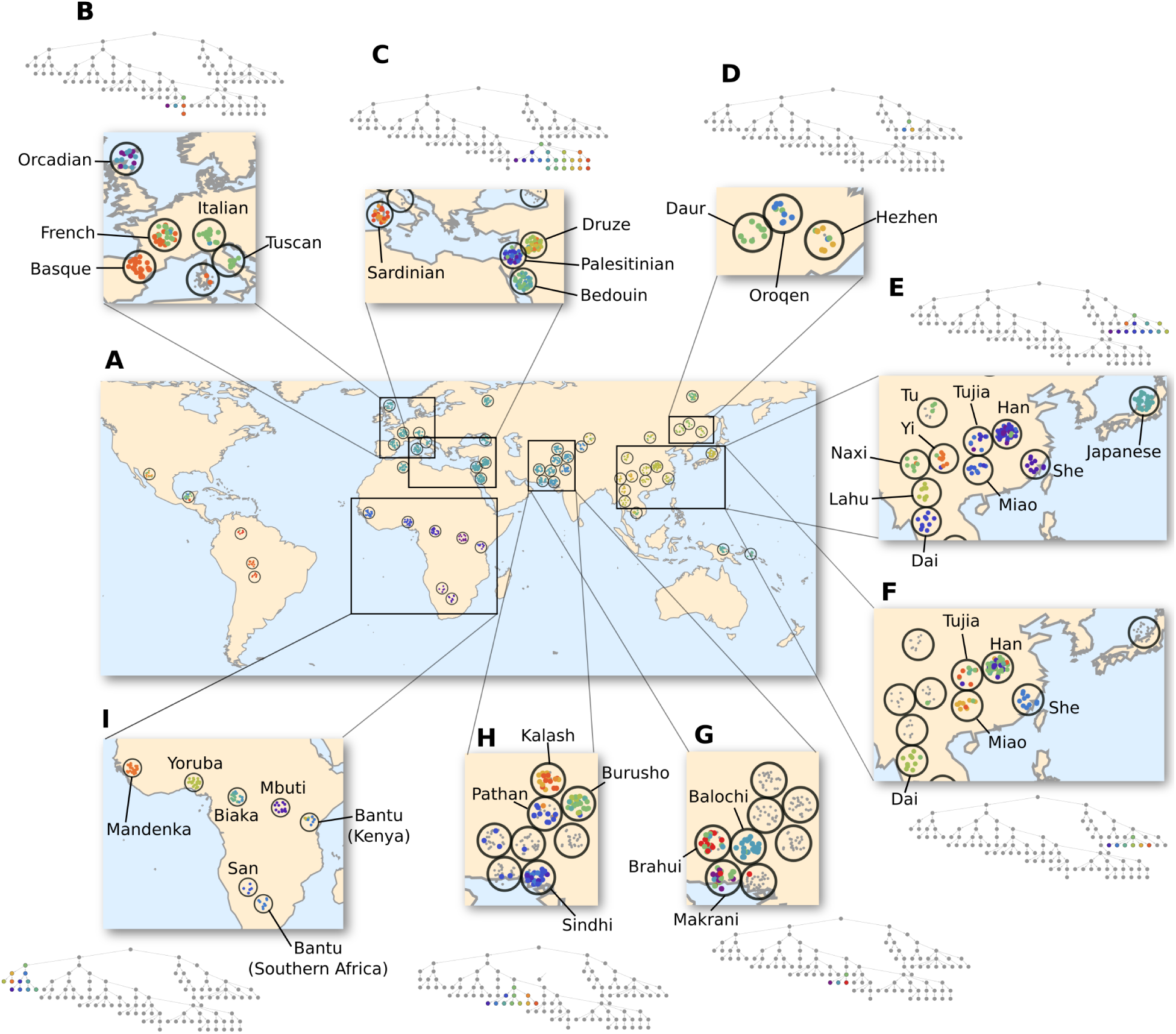
Fine-scale human population structure. Shown is a visualization of the PST on a world map. Each open circle corresponds to one of the 52 groups of the HGDP dataset; group coordinates from Rosenberg (2011) were used, but adjusted such that circles do not overlap (see Fig. S3 for group labels). Individuals were positioned randomly within their corresponding circles, and colored according to the finest-scale cluster to which they were assigned. To illustrate fine-scale structure at a local level, a variety of regions are shown in detail. (B) Branch corresponding to Europe. (C) Branch corresponding to the Mediterranean region. (D) Branch corresponding to northern China. (E) Branch corresponding to Japan and central and southern China. (F) Branch corresponding to central and southern China. (G) Branch corresponding to Balochi, Brahui, and Makrani groups. (H) Branch corresponding to Buruso, Kalash, Pathan, and Sindhi groups. (I) Branch corresponding to Sub-Saharan Africa. In each inset, the branch has been re-colored according to an automatic coloring scheme, which assigns closer colors to clusters positioned closer in the PST, except in (B) and (G), where each cluster was assigned a color manually, irrespective of positioning in the PST.

Once the clusters are assigned colors, each individual is assigned a color corresponding to the finest-scale cluster to which it is assigned. On a geographic map of the region of interest, each individual is placed in its geographic sampling position and colored with its assigned color. This step results in a map reflecting the topology of a particular branch of the PST. By selecting different root clusters and repeating the process, different geographic regions and different levels of structure can be emphasized (e.g., Figs. 3 and 5).

### 2.3 Population structure tree in *Arabidopsis thaliana*

Understanding the population structure of the model plant species *A. thaliana* has been of great interest both for explaining patterns of natural variation and for correctly interpreting signatures of selection and association inferred from genome-wide association studies (Sharbel et al. 2000; Jørgensen and Mauricio 2004; Nordborg et al. 2005; Bakker et al. 2006; François et al. 2008; Platt et al. 2010; Alonso-Blanco et al. 2016; Provart et al. 2016; Durvasula et al. 2017; Lee et al. 2017). Within Eurasia, population structure has been characterized in one study by nine clusters — the group of highly differentiated individuals found primarily in the Iberian Peninsula, and eight other clusters, broadly corresponding to North Sweden, South Sweden, the Iberian Peninsula, Asia, Germany, Italy/Balkans/Caucasus, West Europe, and Central Europe (Alonso-Blanco et al. 2016). Recently, evidence provided by samples from Africa has suggested that African populations are distinct from Eurasian populations (Durvasula et al. 2017). In North America, the population has been found to be relatively unstructured (Jørgensen and Mauricio 2004; Hagmann et al. 2015; Alonso-Blanco et al. 2016).

Here, we analyzed worldwide population structure of *A. thaliana* using our network-based method, in order to better understand hierarchical levels and fine-scale structure in this model species. We studied 1,135 individuals (accessions) from Eurasia and North America (Alonso-Blanco et al. 2016), and 79 individuals from Africa (Durvasula et al. 2017), with the combined dataset containing 1,214 individuals and 2,367,560 SNPs.

We detected a highly hierarchical population structure, with 247 (overlapping) clusters overall and 96 fine-scale (non-overlapping) leaf clusters at the tips of the hierarchy (Fig. 2A). The topology of the inferred PST is highly unbalanced, with one main branch consisting of a single cluster (left main branch in Fig. 2A), and the other main branch containing 246 clusters (right main branch in Fig. 2A). All individuals in the single-cluster left main branch are from North America, representing 71% of the North American sampled population (89 of 125). The right main branch consists of two sub-branches, one representing European individuals (Fig. 2B) and the other representing African, Asian, and North Swedish individuals, and some Iberian individuals designated as “relicts” by previous studies (Alonso-Blanco et al. 2016; Durvasula et al. 2017; Lee et al. 2017) (Fig. 2C). Population structure in the primary European branch shows relatively continuous genetic differentiation over space, as evidenced by the fact that geographically close samples are closely positioned in the PST topology (Fig. 2B). In the primarily non-European branch, population structure is more discontinuous geographically, with a large “gap” between the Asian, Iberian, and North Swedish branches (Fig. 2C). However, looking at fine-scale population structure in Africa and Iberia (Fig. 3F-3G) and in Asia (Fig. S1), population structure appears to be more geographically continuous within these smaller branches. Note that in North Sweden, the sampled area was too small to examine geographic patterns of differentiation within the region.

The separation, at a high hierarchical level, between the European branch and the African, Asian, and North Swedish branches could reflect an ancestral split between these groups (Durvasula et al. 2017), but how these spatially distant populations are related, and why a geographic gap exists in the primarily non-European branch, remain to be investigated. Alternatively, the split partitioning these three groups from the European group may reflect strong genetic similarity within the European branch, rather than genetic similarity between these three groups. Therefore, further study of the population in North Sweden, and increased geographic sampling in Africa and Asia, could help expain these patterns (e.g., Hsu et al. 2019). The continuous differentiation, across hierarchical scales, in the European branch accords with past studies of this geographic region (Sharbel et al. 2000; Nordborg et al. 2005; François et al. 2008; Alonso-Blanco et al. 2016).

Structure can also be ascertained at a very fine scale by examining branches close to the tips of the hierarchy. In the Iberian Peninsula, for example, genetic differentiation can be observed at several scales (Fig. 3). At the broader scale, in the western part of the peninsula, genetic differentiation is aligned along a north-south gradient (Fig. 3B), whereas in the northeastern region, genetic differentiation more closely follows an east-west gradient (Fig. 3C). Distinct groups, correlated with geography, can be seen at fine spatial scales, such as along the northeastern coast (Fig. 3E), in the north (Fig. 3B), and in the central region (Fig. 3D). In Morocco and the Iberian Peninsula, for those individuals belonging to the primary non-European branch in Figure 2A, population structure follows a north-south gradient, with clusters corresponding to small and specific regions in Morocco (Fig. 3G) and to larger regions in the Iberian Peninsula (Fig. 3F). In Asia, population structure corresponds to broad regions (e.g., Altai Mountains, Central Asia, Ural Mountains), but finer-scale patterns are also observed within these regions (Fig. S1).

The formation of such small-scale structure might be attributable to small-scale landscape barriers and resistance to gene flow, but also to adaptation of *A. thaliana* to specific local conditions (Li et al. 2010; Fournier-Level et al. 2011; Horton et al. 2012). For example, the branch corresponding to the coastal population in northeastern Spain has been suggested to be locally adapted to high salinity (Busoms et al. 2015). Selective sweeps suggesting significant local adaptation have also been reported for the population in North Sweden (Long et al. 2013; Huber et al. 2014). Such events could have led to decreased fitness of hybrids and decreases in effective migration, intensifying the differentiation of this population. This phenomenon could perhaps contribute to explaining the non-intuitive placement of North Sweden in the PST, separate from most European samples (Fig. 2C).

The single cluster associated with North America represents most of the samples from that region. This lack of structure, which acords with previous studies (Jørgensen and Mauricio 2004; Platt et al. 2010; Hagmann et al. 2015), could result from a founder effect followed by rapid expansion, producing a fairly homogeneous but widespread population (Platt et al. 2010). However, 29% of the North American samples are clustered within the European branch: 20 in a UK/France cluster, 14 in a Central European cluster, and 2 in a South-Sweden/North-Europe cluster (Fig. S2). These samples include 5 of the 6 individuals sampled along the western coast of North America. The clustering of North America individuals in European clusters suggests a history more complex than a single colonization event. Most individuals perhaps descend from a founder population in the eastern United States (Sharbel et al. 2000; Jørgensen and Mauricio 2004), but other migration events, possibly from Western European sources, might also have contributed to the population.

### 2.4 Population structure tree in humans

The Human Genome Diversity Panel (HGDP) includes genomic samples from 52 human populations from across the world (Cann et al. 2002; Cavalli-Sforza 2005). This set of populations is particularly well-suited as an example, because it has served as a test set for a variety of population structure methods (Corander et al. 2004; Corander and Marttinen 2006; Francois et al. 2006; Patterson et al. 2006; Nievergelt et al. 2007; Corander et al. 2008; Hubisz et al. 2009; Shringarpure and Xing 2009; Jombart et al. 2010; Lawson and Falush 2012; Lawson et al. 2012; Pickrell and Pritchard 2012; San Lucas et al. 2012; Loh et al. 2013; Frichot et al. 2014; Raj et al. 2014; Gopalan et al. 2016; Granot et al. 2016; Hao et al. 2016; Hunley et al. 2016; Zheng and Weir 2016). We generated a PST from 938 HGDP individuals, typed at 647,976 SNPs.

The PST consists of 108 clusters overall, and 57 leaf clusters (Fig. 4). The main branches of the PST correspond to continental-level patterns of human groups, such as Sub-Saharan Africa, East Asia, and the Americas (Fig. 4A), recapitulating structure observed with other methods in these samples (Rosenberg et al. 2002; Jakobsson et al. 2008; Li et al. 2008). Several populations, including Cambodians, Melanesians, Papuans, and Yakut, are not placed within any main continental branch (Fig. 4B). These positionings highlight persistent variability in the placement of these populations; the Oceanians and the Yakut population generally emerge as clusters in detailed analyses, but they have variable placement in relation to the largest clusters in analyses with lower clustering resolution.

Two Native American groups, Maya and Pima, are split into two positions in the PST — one within the branch corresponding to the Americas (4 Maya and 8 Pima individuals; Fig. 4B), and one outside any main continental group (14 Maya and 6 Pima individuals; Fig. 4B). This pattern may reflect previous observations of apparent admixture in the Maya sample, with possible genetic contributions from East Asian and European populations (e.g., Rosenberg et al. 2002; Pickrell and Pritchard 2012). The Pima sample has not previously been detected as an intercontinentally admixed group; because our analysis is based on genetic similarity between individuals, and because the Maya and Pima groups are genetically similar, the clustering of 6 Pima individuals outside the main branch of the Americas might reflect their similarity to admixed Maya individuals, rather than admixture in the Pima themselves.

The PST reveals fine-scale structure, which generally agrees with previous analyses, but in some cases provides finer details. Most groups can be associated with a single cluster or a small cluster-group in the PST (Fig. 4B). Detailed previous analyses of the HGDP sample have been able to clearly separate groups particularly in the Americas and the Middle East, and to some extent in sub-Saharan Africa (Jakobsson et al. 2008; Li et al. 2008). These groups are also clearly separated in the PST (Figs. 4B, 5C, and 5I). Fine-scale structure has been more difficult to resolve in Central/South Asia, East Asia, Europe, and in some sub-Saharan African populations. Our network-based approach is able to identify finer-scale structure for many groups in these regions. In Central and Western Africa, Biaka, Mandenka, Mbuti, and Yoruba are each clustered to a different leaf cluster, with each cluster consisting of individuals from a single group (Fig. 4B and Fig. 5I). In Europe, we identified two Orcadian leaf clusters, containing all individuals from that group, and a leaf cluster consisting of a subgroup of Basque individuals (Figs. 4B). At higher hierarchical levels, we identified a Basque and French cluster (Figs. 4B and 5B). In Central/South Asia, we identified leaf clusters corresponding to many of the groups in the HGDP: Balochi, Brahui, Burusho, Hazara, Kalash, and Makrani (Fig. 4B).

Non-leaf clusters also inform population structure, with clusters corresponding to a Hazara and Uygur group; a Pathan and Sindhi group; a Balochi, Brahui, and Makrani group; and a Burusho, Kalash, Pathan, and Sindhi group (Figs. 4B, 5G, and 5H). In East Asia, three northern Chinese groups are separated in the PST, and the Hezhen and Oroqen groups are each assigned a leaf cluster (Figs. 4B and 5D). The Japanese, Lahu, and Yi groups are also clearly separated, and each is assigned a leaf cluster (Figs. 4B and 5E). A cluster of groups from China is separated into 6 clusters: Dai, Miao, She and three clusters containing Han and Tujia individuals (Figs. 4B and 5F).

Because our coloring scheme differentiates between individuals assigned to leaf clusters and those that are not, it aids in observing groups that are not grouped exclusively in a leaf cluster. For example, in the branch corresponding to northern China, 8 of the 9 Daur individuals are assigned to a non-leaf cluster, but not to the leaf clusters below it, whereas the Hezhen and Oroqen individuals are assigned to leaf clusters. Therefore, although the Daur group is not characterized by any exclusive leaf cluster, it can be visually differentiated from the other two populations in Figure 5D. Similarly, by subtracting individuals from leaf clusters, an Italian and Tuscan grouping can be observed in Fig. 5B, and a Bantu (Kenya and Southern Africa) grouping can be observed in Fig. 5I, although individuals from those groups are not a majority in any specific cluster.

### 2.5 Comparing population structure trees

Large genomic datasets provide opportunities to conduct subsampling analyses on the same dataset, for example to investigate population structure inferred from different genomic regions, or to study the effect of different genome sequencing schemes. To interpret such subsampling analyses, a coherent method for comparing population structure outputs is needed. We therefore developed an approach to compare different PSTs derived from the same dataset.

The approach is based on normalized mutual information (NMI), an information-theoretic measure that evaluates, for a pair of variables, the amount of information gained about a variable by observing a second variable. This measure can be adjusted to apply to a set of hierarchical sets of clusters (Lancichinetti et al. 2009), such as PSTs. Given two PSTs derived from the same set of individuals but not necessarily from the same similarity matrix, the measure produces a score that ranges from 0 to 1, where higher scores indicate greater similarity between the PSTs (see *Methods* for details). Notably, two PSTs might differ both in the topological ordering of the clusters in the tree and in the assignment of individuals to clusters, but they are still comparable using the NMI measure.

Our implementation of the NMI measure is flexible in the sense that it is possible to conduct comparisons not only between two PSTs, but also between subsets of the PSTs. This type of comparison enables a focus on comparing certain features of the PSTs. For example, the approach can compare specific branches of interest, or only fine-scale leaf clusters, by considering in the NMI comparisons only specified subgroups of clusters (see *Methods* and Supplementary Information).

### 2.6 Information gain by increased SNP coverage

We estimated the effect of increasing the number of loci used in the inference of PSTs by comparing the PST inferred from an entire dataset with PSTs inferred from subsamples of the dataset, using the information-theoretic NMI measure. This step was accomplished by evaluating NMI between PSTs, one inferred from the full dataset (Figs. 2 and 4), and one inferred from a subsample of the data. We also computed NMI for fine-scale structure by examining only the leaves of the hierarchies. For this purpose, without considering SNP positions, we randomly subsamped all SNPs to produce subsets of 0% to 98% of the original dataset, in increments of 2%. For each frequency value, 100 random subsamples were taken, and from each of these subsamples, a PST was inferred and compared to the full-data PST.

The results (Fig. 6) indicate that information gain is rapid for the first 50,000 SNPs, in both the *A. thaliana* and human datasets. In *A. thaliana*, further SNP coverage results in only modest information gain, suggesting that most population structure is revealed in this dataset using about 100,000 SNPs (Fig. 6A). For the human dataset, more information was gained as SNP coverage continued to increase (Fig. 6B). These results, particularly for the *A. thaliana* dataset, are consistent with previous studies on lower-density genetic data that have shown that the resolution of population structure detected tends to saturate when enough loci are included in the analysis (e.g., Manel et al. 2002; Turakulov and Easteal 2003; Rosenberg et al. 2005; Morin et al. 2009; Haasl and Payseur 2011; Bryc et al. 2013); however, for the human dataset, information gain is not saturated even with several hundred-thousand SNPs. In both datasets, with increased SNP coverage, more information is gained regarding fine-scale structure than about coarse-scale structure. For example, increasing the number of SNPs from 10% of the dataset (236,756 SNPs for *A. thaliana* and 64,797 SNPs for humans) to 90% SNP coverage (2,130,804 SNPs for *A. thaliana* and 583,173 SNPs for humans), increases NMI by 3.7% for *A. thaliana* and 23.7% for humans when considering the entire PST, whereas the increase is larger when considering the finest-scale clustering at the PST leaves, with NMI increases of 11.1% for *A. thaliana* and 77.1% for humans (Fig. 6).

**Figure 6:**
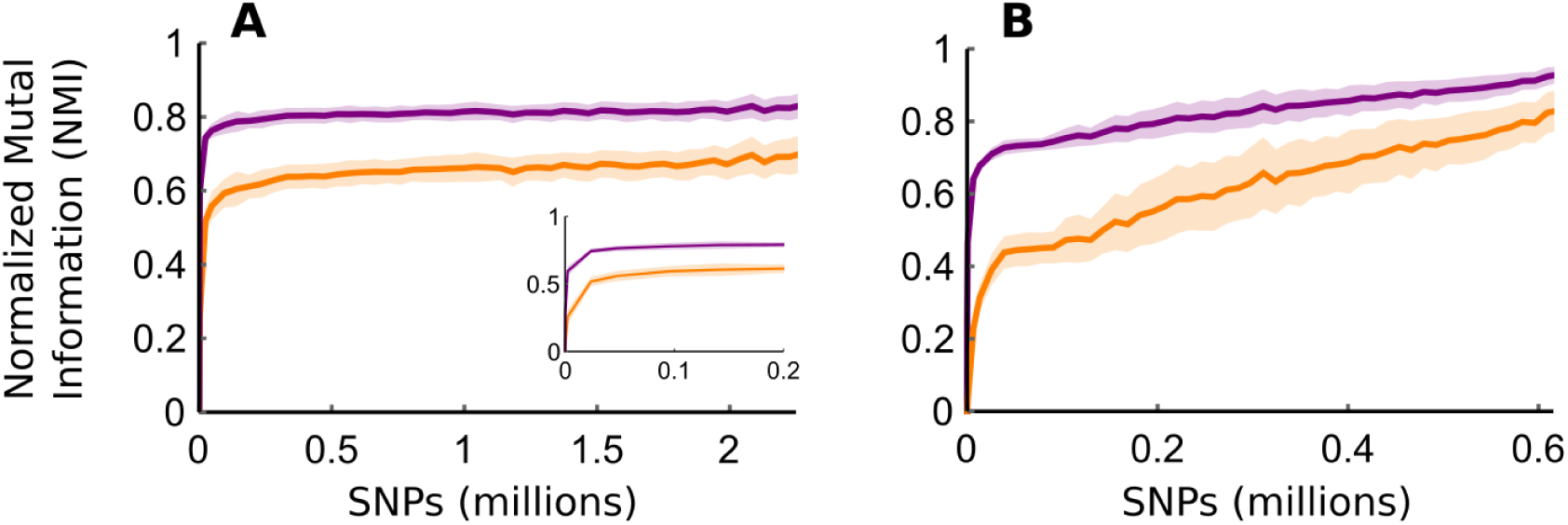
Normalized mutual information (NMI) between the PST inferred using the entire genome and PSTs inferred from subsampled fractions of the genome. The NMI values evaluate the amount of information gained on hierarchical population structure by sampling a fraction of the dataset. The mean NMI across 100 random subsamples for each SNP coverage value considering the entire PST topology is shown in purple; mean NMI considering only the finest-scale leaf clusters is shown in orange. Shaded regions show standard deviations across 100 sampling replicates. (A) *A. thaliana* dataset. Inset shows NMI for subsamples below 200,000 SNPS. (B) Human dataset. NMI values saturate at values below 1 because cluster assignments often switch at fine scales (e.g., between PST leaves) when PSTs are inferred from subsampled data.

## 3 Discussion

### 3.1 Population structure inference using networks

The evolutionary processes shaping genetic variation are often complex, occurring at multiple spatial levels and over multiple time periods. Network-based methods are well-suited for population structure inference because they enable a natural hierarchical perspective. Our approach is data-driven and does not require assumptions or pre-specified models regarding the biological processes responsible for shaping the genetic patterns. The visualization technique presented here, with PST diagrams and corresponding map colorings, is useful for summarizing information on population structure while preserving information at both broad and fine scales.

We demonstrated the applicability of the NMI measure for comparing PSTs to understand the information gained by an increased number of genetic markers. This measure can also be useful for other types of analyses, such as for comparing PSTs inferred from different genomic regions or different types of genetic markers.

### 3.2 Comparison to other approaches

In Table 1, we summarize the features of our network approach in relation to those of popular classes of population structure inference approaches.

**Table 1:** Comparison of features of selected population structure inference approaches representing the major families of methods.

| Inference approach | Reference | Putative pre-definition of groups used in analysis? | Clustering of individuals into groups integral to method? | Hierarchical representation of population structure integral to method? | Requires pre-definition of biological model? |
| --- | --- | --- | --- | --- | --- |
| F-statistics (e.g., Fst) | Excoffier et al. 1992<br>Holsinger and Weir 2009 | Yes | No | Yes | Yes |
| Dimension reduction visualization (e.g., PCA, MDS, UMAP) | Menozzi et al. 1978<br>Jombart et al. 2009 | No | No, but possible with post-analysis methods, e.g. K-means | No | No |
| Tree-based (e.g., TreeMix) | Pickrell and Pritchard 2012 | Yes | No | Yes | Yes |
| Tree-based (e.g., Neighbor-joining) | Bowcock et al. 1994 | No | No | Yes | No |
| Model-based (e.g., STRUCTURE, ADMIXTURE) | Pritchard et al. 2000<br>Alexander et al. 2009 | No (optional) | Yes | No, but multiple models with different numbers of clusters can inform hierarchical structure | Yes |
| Network-based (NetStruct_Hierarchy) | This paper | No | Yes | Yes | No |

The network-based approach, unlike F-statistics or some commonly-used tree-based methods, does not require an assumption of putative populations, and it clusters individuals into groups in a data-driven manner. Unlike in multivariate analysis methods such as PCA, in the network method, clustering is naturally conducted using community-detection procedures. Unlike in STRUCTURE-like methods, the PST generated by the network method depicts hierarchical levels of structure without the need to predefine the number of clusters or to run the analysis at different values for this quantity. Detection of multiple hierarchical levels of structure is integral to the network approach, without assumptions on the number of clusters, the hierarchical topology of population structure, or putative groupings of individuals; all these characteristics are generated from the data. Such features of the method are beneficial particularly when little information is available for specifying models or for defining putative groups, and when the hierarchical level of interest is unclear.

We demonstrated the features of our approach in two datasets. In our application to *A. thaliana* and humans, the PSTs distinguish between groups on a fine scale. The method uncovers clusters that have not been straightforward to distinguish in PCA or STRUCTURE-like analyses in previous studies (Li et al. 2008; Durvasula et al. 2017).

Ideally, inference of population structure should be conducted using more than one approach, to demonstrate robustness and to capitalize on the different features of the different approaches (Table 1). For example, in some cases, we observe in a PST a split between a large group and several smaller groups. These splits are sometimes difficult to interpret, because it is unclear whether they represent genetic similarity between the smaller groups, or strong genetic similarity between individuals in the larger group. For example, the *A. thaliana* split between a branch with African/Asian/North Sweden groups and a large European branch (Fig. 2), and the human split between the Oceanian/Cambodian/Yakut/North American groups and a large mainly-Eurasian branch (Fig. 4A), might be driven by genetic similarities between the groups in the same finer-scale branch, or by strong between-individual genetic similarities in the coarser branch. The distinction of these two scenarios can be examined by leveraging advantages of different classes of methods. For example, once clusters and groups of interest have been ascertained by an approach that does not require definition of putative populations, genetic similarities between the identified groups can be computed using methods that rely on putative group assignments (Table 1); network methods can also aid in investigating the cohesiveness of different clusters, for example using strength-of-association-distribution (SAD) analysis (Greenbaum et al. 2016).

### 3.3 Conclusions

We have presented a network-based approach for revealing population structure at multiple hierarchical levels. This structure is summarized in a novel form, the PST, and can be visualized on geographic maps. The method is applicable to whole-genome data. The approach is data-driven and computationally efficient, allowing analysis of large genomic datasets. Our method for comparing PSTs using the NMI measure is potentially useful for conducting studies that investigate population structure inferred from subsamples of the data.

## 4 Acknowledgments

We thank Arun Durvasula, Andrea Fulgione, and Angela Hancock for help with the *A. thaliana* data, and Jonathan Kang for help with HGDP dataset. We also thank Ellie E. Armstrong, Danny Hendler, and Stefan Prost for helpful discussions. This study was supported by NIH grant R01HG005855 awarded to NAR.

## 5 Author contributions

GG conceived the study and drafted the manuscript. GG and AR developed and implemented the approach, and analyzed the *A. thaliana* and human datasets. ART and NAR contributed ideas to the framing and implementation of the project. All authors read and commented on the manuscript.

## 6 Data availability

The code for running the analyses (called NetStruct Hierarchy) can be found in https://github.com/amirubin87/NetStruct_Hierarchy along with a user manual and a tool for plotting maps and coloring the clusters. The *A. thaliana* genetic dataset was extracted from www.mpipz.mpg.de/hancock/downloads, which stores a unified datast of the 1001 Genomes database (also found at `1001genomes.org`) and the data from Durvasula et al. (Durvasula et al. 2017). Geographic locations of the *A. thaliana* samples were extracted from the 1001 Genomes database at `1001genomes.org` and from Durvasula et al. (Durvasula et al. 2017). The HGDP data was downloaded from the HGDP website hagsc.org/hgdp.

## 7 Methods

### 7.1 Constructing genetic-similarity networks

The first step of network-based population structure detection is to use genomic data to generate a genetic similarity network, with individuals treated as nodes and interindividual genetic similarities as edges (Fig. 1). Here, we use a frequency-weighted allele-sharing similarity measure. Allele-sharing measures count the number of shared alleles between each pair of individuals over all loci. Because individuals that share rare alleles are more likely to belong to the same subpopulation than are those that share common alleles, we assign more weight to shared rare alleles than to shared common alleles. There are different ways to design weighting schemes to generate genetic similarity networks (Greenbaum et al. 2016); here we adopt a linear weighting scheme for its simplicity, but this weighting scheme can in principle be replaced by other weighting schemes if there is reason to tailor weights such that different weights are assigned to different allele frequency classes.

Considering a locus *ℓ*, for individual *i* with alleles *a* and *b* and individual *j* with alleles *c* and *d*, the frequency-weighted allele-sharing similarity is defined as (Greenbaum et al. 2016):

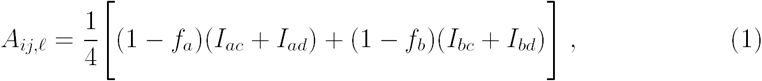

where *I*_*xy*_ = 1 if *x* and *y* are identical and *I*_*xy*_ = 0 otherwise, and *f*_*a*_ and *f*_*b*_ are the frequencies of alleles *a* and *b* in the entire sample. The genetic-similarity edge connecting individuals *i* and *j* is weighted by the mean genetic similarity over all *L* loci available for this pair of individuals, 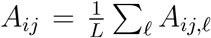. When some data are missing, *A*_*ij*_ is computed based on all those loci for which both individuals *i* and *j* are scored. This measure is symmetric (*A*_*ij*_ = *A*_*ji*_) and defines an inter-individual genetic-similarity network, where *A*_*ij*_ is the adjacency matrix of the network.

### 7.2 Community detection

By a subpopulation in a structured population, we mean a group of individuals that share a common genetic history, and that are therefore more related to one another on average than they are to individuals outside the group. Individuals are then expected to share more genetic similarity edges within subpopulations than between subpopulations. In network science, groups of nodes that are strongly connected within the group, forming dense subnetworks, are termed “communities” (Newman 2002). By detecting the community structure of genetic similarity networks, the genetic population structure can be revealed (Greenbaum et al. 2016).

Many algorithms for community detection have been developed, including for weighted networks for which weights are assigned to the edges (Fortunato 2010; Yang et al. 2016). Many of these methods are based on a quality function that evaluates community partitions, and that is termed “modularity” (Newman 2006). This function, given a network and a partition of the network, returns a value between −1 and 1. Modularity is computed by analytically comparing the intra-community densities of a partition of the network with the expectation of intra-community densities over an implicit distribution of “random networks” that preserve the node weighted-degrees (Newman 2006; Eq. 1 in Greenbaum et al. (2016)). Therefore, high modularity scores reflect a community partition in which within-community connections are more frequent and more highly-weighted than would be expected in a random network with the same degree distribution. Many community-detection algorithms aim to efficiently identify a community partition that maximizes the modularity score. Here, we use a particularly time-efficient weighted community-detection method, the Louvain method (Blondel et al. 2008), which attempts to maximize modularity through a greedy search using a deterministic iterative process.

### 7.3 Edge-pruning

In structured populations, genetic variants are more likely to be shared within sub-populations than between subpopulations. Different genetic processes might have occurred over different spatial scales, either at different points in time or simultaneously, and therefore, the genetic signature is a complex aggregate of structure at several scales. Fine-scale population structure is characterized by groups of individuals that are strongly related, and that therefore are expected to be represented in subnetworks that contain only the strongest genetic similarity edges. Coarse-scale population structure, on the other hand, is characterized by both strong and weak genetic similarities.

In order to detect population structure at multiple hierarchical levels, we perform an iterative edge-pruning process, where we sequentially remove edges below a genetic-similarity threshold. Here, we start at threshold 0 (i.e. all edges are included), and calculate the community structure. If only a single community is detected (no community structure), then we increment the threshold, remove the edges below the threshold, and again apply community detection. Once community structure is detected, the network is split, and the process is repeated for each community independently. This process of subdividing communities forms a PST, where clusters closer to the root represent coarse-scale structure and clusters at the leaves represent fine-scale structure.

We selected a threshold increment of 10^−4^ for the analysis of the *A. thaliana* dataset and 10^−3^ for the human dataset, such that the final output, the PST, does not substantially change with each increment. For other datasets, we recommend inspecting the typical differences between the sorted weights of the edges, selecting a threshold larger than the typical gaps between consecutive edge weights, and testing several threshold increments, at different orders of magnitudes, to identify stable outputs. Higher threshold increments are expected to result in fewer hierarchical levels in the PST, because removal of many edges between applications of the community-detection procedure may result in hierarchical levels being skipped over.

The PST generated in this way is entirely data-driven, with the number of fine-scale and coarse-scale clusters determined only from the genetic data. Note that the population structure could include small clusters containing relatively few individuals. In some cases, even clusters containing only a few individuals might be informative, because they might represent sparsely sampled population fragments or familial groups. In other cases, such small groups may not be of interest. We therefore impose a condition on the process by specifying the minimum number of individuals in a cluster, where communities below that size are not considered clusters and do not induce a split in the PST. Because large genomic datasets can potentially generate a fine-scale level of structure, we kept these minimum sizes small, and ignored clusters smaller than five individuals for both datasets. In other datasets, keeping small clusters may reveal fine-scale structure, but this high-resolution structure could be less reliable due to potentially non-representative individuals; we recommend testing several minimum cluster sizes, and keeping in mind the minimal group sizes for which meaningful interpretation can be given (e.g., the smallest predefined group in the human dataset includes 5 individuals).

### 7.4 Generating geographical PST maps

We use a coloring scheme to visualize population structure. This scheme represents fine-scale structure while accounting for the hierarchical topology of population structure, and it allows identification of relevant differentiation patterns (Figs. 2–5). For each coloring, we select a root cluster from the PST. This cluster defines the scale and focus of the coloring; the coloring is intended to visualize the branch leading from this root cluster to the leaves.

We start with an interval [0, 1] representing a color gradient. Here we use the “Rainbow” color gradient as implemented in Mathematica (Wolfram Research and Wolfram Research 2018), with each number in the interval representing a color on the gradient; see Fig. S3 for an explicit representation of this color scheme function. We assign the starting interval to the root cluster. We then evenly split the interval among all immediate daughter clusters of the selected root cluster. We repeat this process, where in each step, a sub-interval of the parental interval is inherited by daughter clusters. In other words, if a cluster *C* is assigned the interval [*a, b*], and it has *n* daughter clusters *C*_1_,…,*C*_*n*_, then the daughter cluster *C*_*i*_ is assigned the interval 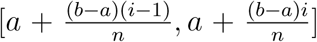. This method assigns smaller intervals to clusters as we advance toward the leaves.

The process is terminated when leaves have been reached on all sub-branches. We then assign a single color to each cluster, by taking the color associated with the midpoint of the interval assigned to that cluster. In other words, if cluster *C* is assigned the interval [*d, e*], then *C* is assigned the color corresponding to 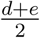. In this way, fine-scale clusters that are far from the root cluster and that share a common parent cluster one level above will tend to be assigned “closer” colors than will clusters that share a parent cluster nearer to the root cluster.

Each individual can be assigned to multiple clusters in different hierarchical levels, and we assign a color to an individual based on the finest-scale cluster to which it is assigned (i.e. the cluster closest to a leaf). This color assignment of individuals is then used for plotting geographic maps, where for each individual, a point is plotted at its sampling location, colored in its assigned color.

In this way, the many fine-scale clusters that are detected in large genomic datasets can be presented simultaneously, while allowing the broader genetic patterns to be clear on each map and at each scale by the coloring scheme. For example, although Figures 2 and 3 use many colors, some of which are visually indistinguishable, the broad patterns of genetic differentiation can be easily comprehended. When structure at a fine scale is of interest, the coloring scheme can be repeated while selecting different root clusters, thus focusing on different sub-branches of the PST. This method makes it possible to rescale the coloring scheme to the appropriate geographic scale, and finer-resolution genetic differentiation patterns become apparent (e.g. Fig. 3). A caveat of our coloring scheme is that when the number of daughter nodes is odd, one of the daughter nodes and the parent node are assigned the same color; this may limit visual interpretation, particularly when a parent cluster and its daughter clusters are visualized near the leaves of the tree. In such cases, we suggest assigning colors manually (e.g., Figs. 5B and 5G). It is important to note that the PST holds information on all hierarchical levels; the coloring strategy we suggest only aids with visualizing the relationship of genetic differentiation patterns to geography.

All maps were plotted using Mathematica (Wolfram Research and Wolfram Research 2018) GeoGraphics, which imports maps from Wolfram | Alpha (Wolfram Alpha 2018).

### 7.5 Pre-processing of the datasets

For *A. thaliana*, we used data from the 1001 Genomes project, which includes whole-genome SNP data of 1,135 *A. thaliana* diploid individuals (accessions) from Eurasia and North America (Alonso-Blanco et al. 2016), and data from Durvasula et al. (2017) on 79 individuals from Africa. For the combined dataset of 1,214 individuals, we considered only biallelic SNPs, and filtering included removal of all SNPs with more than 10% missing data and all SNPs with minor allele frequencies < 0.01. After filtering, the combined dataset included 1,214 individuals and 2,367,560 SNPs.

For humans, we used 938 individuals from the HGDP dataset. This subset of individuals contain those that remain after removing low-quality samples, replicated samples, and closely-related individuals (Rosenberg 2006; Li et al. 2008). We used the same data filtering as for the *A. thaliana* dataset for the purpose of tractability, i.e., only biallelic SNPs were considered, and SNPs with more than 10% missing data were removed, as were loci with minor allele frequencies < 0.01. After filtering, the dataset included 938 individuals and 647,976 SNPs (of the 660,918 SNPs in the initial HGDP dataset).

### 7.6 Comparing PSTs using NMI

To compare two PSTs derived from the same set of individuals *I*, we compute the normalized mutual information (NMI) between representations of the two PSTs as sets of partitions of *I* (Lancichinetti et al. 2009; McDaid et al. 2013). NMI quantifies the amount of information shared by two random variables, or the amount of uncertainty on one variable that is reduced by observing the other variable, as measured in bits.

In order to apply this measure, we represent a PST, *X*, with *k* clusters, as a partition *X*_1_,…,*X*_*k*_ of the set of individuals *I* (*X*_1_ ∪ *X*_2_ ∪ …∪*X*_*k*_ = *I*). The sets *X*_1_, …, *X*_*k*_ need not be mutually disjoint, and indeed in PSTs, clusters can be nested within other clusters (e.g., a daughter cluster is nested in the parent cluster). The NMI measure computes, for two PSTs *X* and *Y* represented as partitions, a score between 0 and 1, where higher scores represent higher similarities between the PSTs. We summarize the computation in Supplementary Information, and refer the readers to Lancichinetti *et al*. (2009) and McDaid *et al*. (2013) for the full details of this computation.

In some cases, it is of interest to compute NMI not for the entire PST, but rather for a subset of the clusters. For example, we computed NMI for leaf clusters of PSTs in order to compare the fine-scale structure described by the PSTs. Given two PSTs, *X* and *Y*, represented as partitions *X*_1_, …,*X*_*k*_ and *Y*_1_, …,*Y*_*m*_, this computation is performed by considering only a subset of the sets in the partition, and computing NMI using only these subsets. However, because NMI requires both PSTs to be defined on the same set of individuals, a problem arises when the unions of the subsets of clusters are different. In other words, if we consider a subset of *k*′ ≤ *k* clusters from PST *X* and a subset of *m*′ ≤ *m* clusters from PST *Y*, we might have 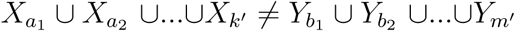. This problem can be addressed by removing or adding individuals to clusters in a standardized way, so that the unions of the subsets of clusters become equal; our analysis of information gain by increased genomic density provides an example of how such a case can be addressed.

### 7.7 Information gain by increased sequencing

The NMI measure quantifies the information shared between two PSTs. If one of the PSTs considered was ascertained using strictly lower-density genetic data, then NMI can be used to evaluate the information gained by increasing the genomic density of the data, as we have done here for the *A. thaliana* and human datasets.

We calculated NMI for PSTs constructed from a subsample of the genome to simulate different fractions of the genome sequenced (0% to 100% in intervals of 2% of the number of SNPs in the dataset). First, the data were randomly partitioned into 10,000 approximately evenly-sized groups of SNPs, subsampled randomly from all SNPs without replacement, regardless of position in the genome. For each fraction of the genome, *f*, 100 random subsamples of the genome, each summing up to a fraction *f* of the genome, were taken without replacement (constructed using the SNP groups, for computational efficiency). For each such subsample, the PST was derived, and the NMI between that PST and the PST derived from the entire genome (Fig. 2A) was computed. The mean NMI and standard deviations for each *f* were then calculated.

In addition, we also considered the information gained when considering only fine-scale structure. This step used a similar process, only that the NMI was calculated by considering only the leaf clusters of the PSTs. Because the union of the leaf clusters does not necessarily cover the entire population, due to some individuals possibly not being assigned to any leaf, each individual not assigned to any leaf was added to all the leaves in the branch below the finest-scale cluster to which that individual was assigned. In other words, for PSTs *X* and *Y*, with leaf clusters *X*_1_,…,*X*_*r*_ and *Y*_1_, …,*Y*_*s*_, respectively, we added each individual *d* in 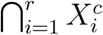 (where 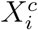 is the complement of *X*_*i*_) to all leaf clusters that belong to the branch emerging from the cluster ⋂_*x*∈*X*|*d*∈*x*_ *x*; we do the same for all individuals in 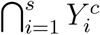 (see Supplementary Information for an illustrative example). We compute NMI for the sets *X*_1_,…,*X*_*r*_ and *Y*_1_, …,*Y*_*s*_ with the added individuals, which now cover the full set of individuals. This procedure generates a comparison between the finest-scale structures described by the PSTs.

### 7.8 Computational efficiency

A major issue with analyses of large genomic datasets is computational efficiency. The method we present requires two stages: (1) construction of the genetic similarity network, and (2) construction of the PST. The first stage amounts to construction of a pairwise matrix of size *n*^2^, where *n* is the number of individuals. Each element requires *ℓ* calculations, where *ℓ* is the number of loci. To apply Eq. 1, calculation of allele frequencies for all loci is required. The computation time for the first stage is therefore *O* (*ℓn*^2^ + *ℓn*) = *O*(*ℓn*^2^) time. This computation time can be reduced substantially by computing the elements of the matrix for each of the loci in parallel: given *c* computation threads, the computation time can be reduced to 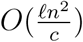 time.

The computation time of the second stage depends on the topology of the PST. With the community-detection Louvain algorithm, each community detection computation takes *O i* log *i* time, where *i* is the number of individuals in the network examined (Blondel et al. 2008). Therefore, for each level of the hierarchy, assuming we have *k* clusters in that level, where cluster *j* has size *i*_*j*_, the computation time is 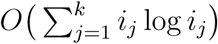 for that hierarchical level. Because 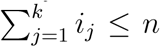 and *i*_*j*_ ≤ *n* (and therefore log *i*_*j*_ ≤ log *n*) for any *j*, the run time of each level is bounded above by *O*(*n* log *n*). For a PST with *d* levels, the overall computational time of this step is, therefore, smaller than *O dn* log *n*. The factor *d* is not known *a priori*, and could differ for different datasets; *d* is bounded by *d* ≤ *n*, although in practice it would typically be much smaller.

In practice, for whole-genome data, the first step (constructing the matrix) takes substantially more time to compute than the second stage (iterative edge-pruning and community detection); the second step does not depend on the number of loci in the data. Using a single computation thread, the first step would require approximately 82 days for the *A. thaliana* dataset and 14 days for the human dataset. We used 1,000 computation threads, and the computation took 2 hours for the *A. thaliana* dataset and 20 minutes for the human dataset. The second step, using a single computation thread, required 49 minutes for the *A. thliana* dataset and 1.5 minutes for the human dataset.

