## Supplementary Materials for "Network-based hierarchical population structure analysis for large genomic datasets"

### Supplementary Information

- Appendix A — *A. thaliana*
- Appendix B — Humans
- Appendix C — Color scheme
- Appendix D — Comparing PSTs using the NMI measure

#### Appendix A — *A. thaliana*

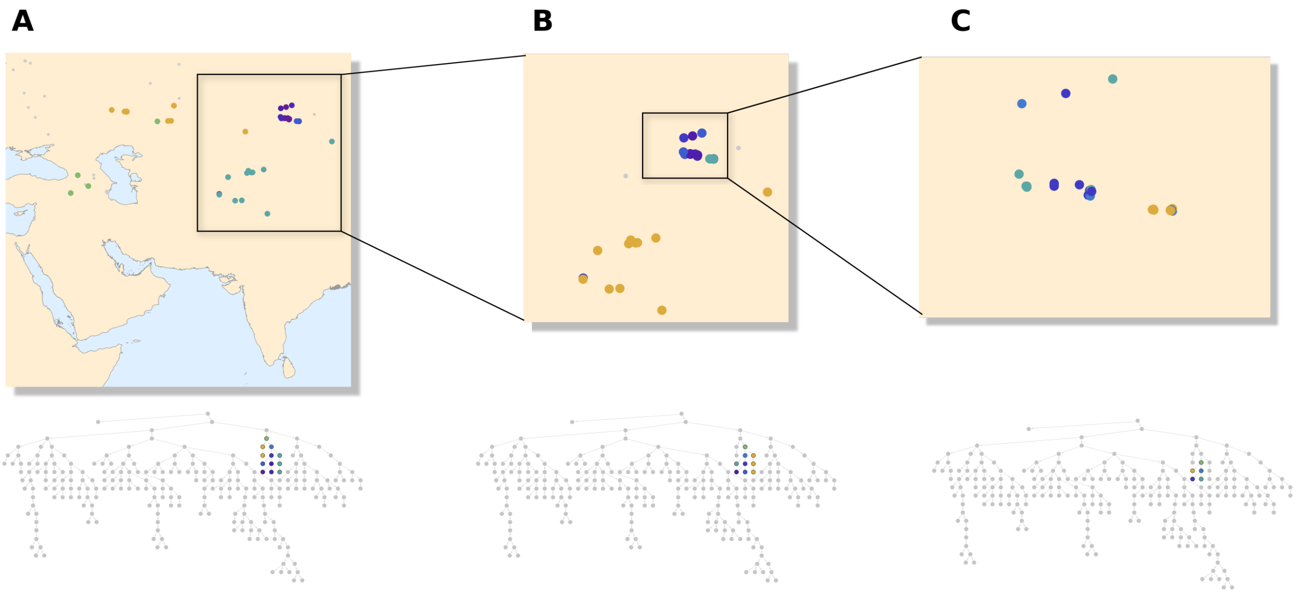


**Figure S1:** ***A. thaliana*** **population structure in Asia.** On each map, a branch of the inferred PST from Figure 2 is visualized, and below each map is the PST colored in the same manner as in the map. (A) The branch corresponding to Asia (purple branch in Figure 2A). (B) A branch corresponding to Central Asia and the Altai Mountains, showing genetic differentiation between these two geographic regions. (C) Fine-scale population structure in the Altai Mountains.

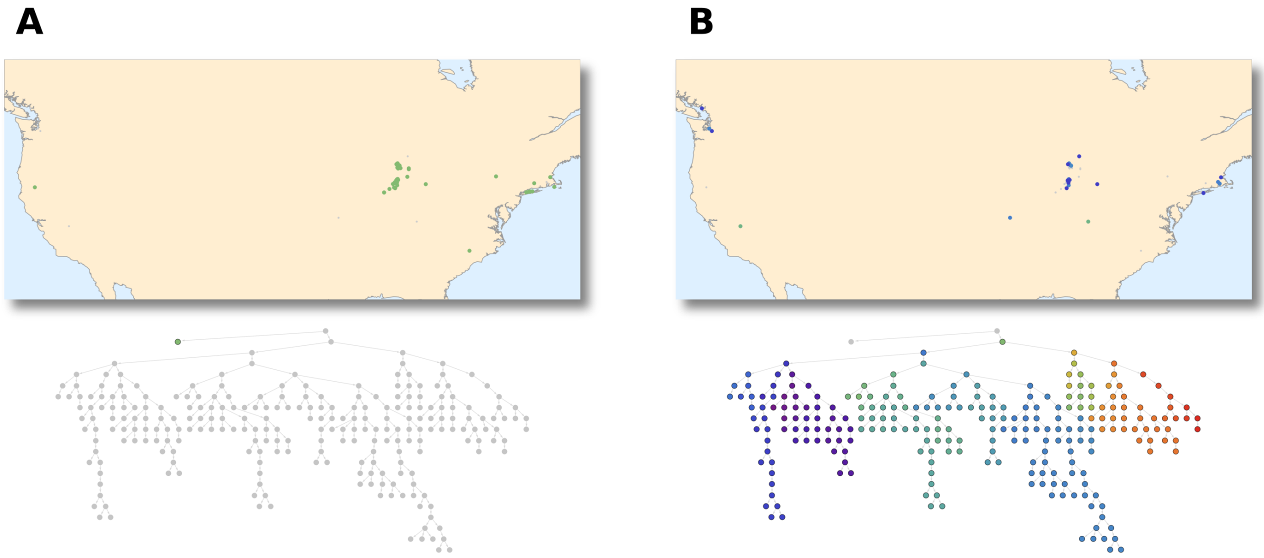


**Figure S2: *A. thaliana* population structure in North America**. On each map, a branch of the inferred PST from Figure 2 is visualized, and below each map is the PST colored in the same manner as in the map. (A) The main left branch of the inferred PST (pink branch in Figure 2A), with a single cluster found only in North America. Of the 125 samples from North America, 89 were assigned to this branch. (B) The 36 individuals in North America that were assigned to European clusters (blue branch in Figure 2A).

### Appendix B — Humans

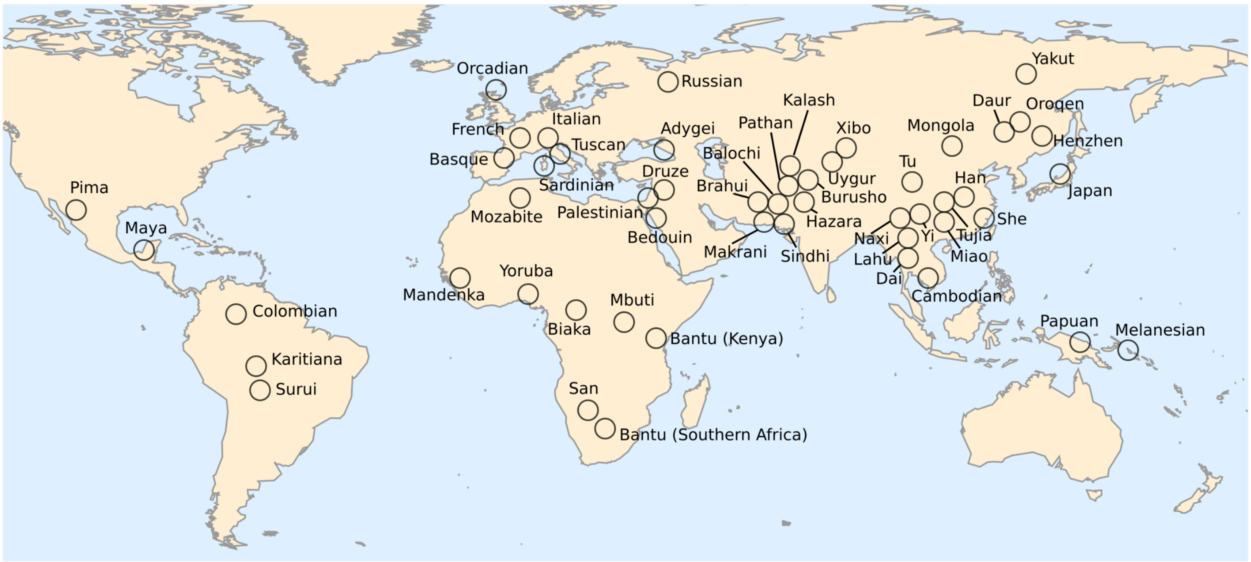


**Figure S3: Population group names and geographic positions for the human dataset**. The map shown corresponds to the map in Fig 4B, with each of the 52 population groups of the HGDP dataset labeled.

### Appendix C — Color scheme

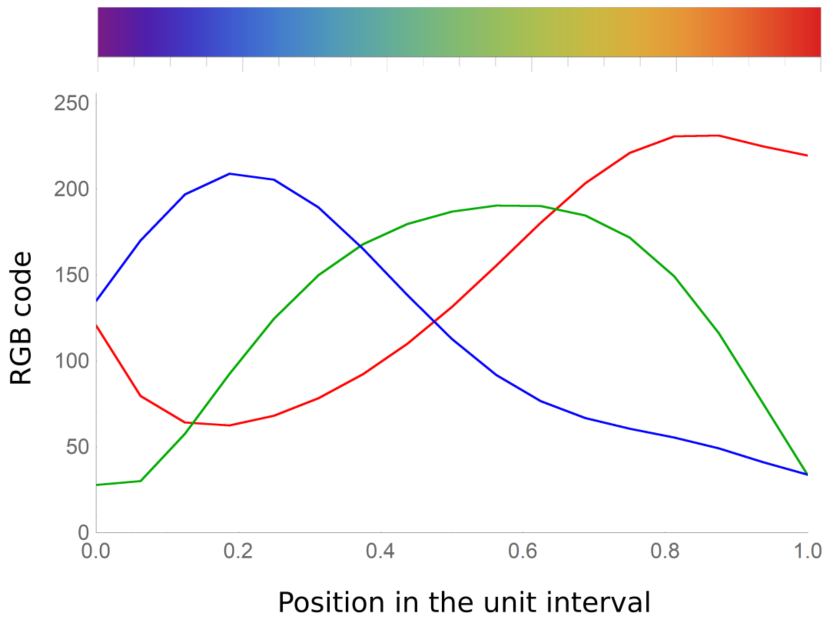


**Figure S4: The RGB codes of the color scheme used for visualization**. The RGB code (with values between 0 to 255) for the three base colors (red, green and blue curves) is shown as a function of a number in the interval [0,1], as implemented in the “Rainbow” color scheme function in Mathematica. At the top are the colors corresponding to the associated RGB code. To generate this plot, colors were sampled at intervals of 0.000001 on the x-axis.

### Appendix D - Comparing PSTs using the NMI measure

#### Computing NMI for PSTs

To compare two PSTs derived from the same set of individuals $I$, we compute the normalized mutual information (NMI) between representations of the PSTs as partitions of $I$ [1]. We follow the NMI definition of McDaid *et al.* [2] ($NMI_{LFK}$ in their work), which differs slightly from the original definition of Lancichinetti *et al.* [1].

For two discrete random variables $X$ and $Y$, their mutual information, MI, is defined as [3]:

$$MI(X,Y) = H(X) - H(X|Y) = H(Y) - H(Y|X) \ , \tag{S1}$$

where $H(X)$ and $H(Y)$ are the marginal information entropies of $X$ and $Y$, and $H(X|Y)$ and $H(Y|X)$ are the conditional information entropies.

In the context of the information content of clusters of data, MI is defined using representations of the clusters as partitions of the data induced by the clustering [1, 4]. We represent a PST, $X$, with $k$ clusters, as a partition $X_1$,...,$X_k$ of the set of individuals $I$ of size $n$ ($X_1 \cup X_2 \cup ... \cup X_k = I$; $|I| = n$). The sets $X_1$,...,$X_k$ are not necessarily mutually disjoint, since some clusters might be nested within other clusters in the topology of the PST. The information entropy of a PST $X$ is defined as [4] (equation below Eq. 4 in [2]):

$$H(X) = \sum_{i=1}^{k} \left[ -|X_i| \log_2 \frac{|X_i|}{n} - |X_i^c| \log_2 \frac{|X_i^c|}{n} \right], \tag{S2}$$

where $|X_i|$ is the size of cluster $X_i$, and $X_i^c$ is the complement of cluster $X_i$. The entropy of $X$ (with $k$ clusters) conditioned on PST $Y$ (with $m$ clusters) is defined as (Eqs. 3 and 4 in [2]):

$$H(X|Y) = \sum_{i=1}^{k} \min_{j=1,...,m} H(X_i|Y_j). \tag{S3}$$

Here, we follow McDaid *et al.*, and define $H(X_i|Y_j)$ as (Eq. 1 in [2]):

$$H(X_i|Y_j) = -a \log_2 \frac{a}{n} - b \log_2 \frac{b}{n} - c \log_2 \frac{c}{n} - d \log_2 \frac{d}{n} + (b+d) \log_2 \frac{b+d}{n} + (a+c) \log_2 \frac{a+c}{n}, \tag{S4}$$

with $a = |X_i^c \cap Y_j^c|$, $b = |X_i \cap Y_j^c|$, $c = |X_i^c \cap Y_j|$, $d = |X_i \cap Y_j|$. Eq. S4 quantifies the lack of information between the clusters. If $x$ and $y$ are two identical clusters, then $b = c = 0$ and there is no lack of information between the cluster, i.e. $H(x|y) = 0$.

The minimum entropy taken in Eq. S3 is intended to associate each cluster in $X$ with its most similar cluster in $Y$, in terms of information content. In other words, each cluster in $X$ is compared with a single cluster in $Y$, the one for which its conditional entropy is minimal.

One possible complication of this procedure can arise when the minimal entropy is attained for two clusters that are almost complementary, which are dissimilar in the individuals assigned to them, yet are similar in their information content regarding clustering assignment. For example, if $x$ is a cluster in $X$, and $y = x^c$ is a cluster in $Y$, then $\min_j H(x|Y_j)$ is minimized by $y$ (since $H(x|y) = 0$), although $x$ and $y$ do not share any individuals. To avoid this possibility, an additional restriction is applied in Eq. S3, to allow only those clusters in $Y$ that are far from being the complement of a cluster $x$ in $X$ to be considered as those minimizing the conditional entropy of $x$; see Lancichinetti *et al.* [1] for details on this constraint.

Although Eq. S1 implies that MI is symmetric, if we were to apply Eq. S1 with Eqs. S2 and S3, we would not necessarily get a symmetric measure, because it is not guaranteed that if a cluster $x$ in $X$ is associated with a cluster $y$ in $Y$ (i.e. $H(x|Y_j)$ is minimized by $y$), then $y$ will be associated with $x$ (since perhaps $H(y|X_i)$ is not minimized by $x$). Additionally, the values of MI defined using Eq. C3 would depend on the numbers of clusters in the PSTs, and would not be comparable with MI measurements between other PSTs. Therefore, a normalization of MI is performed, to ensure that the values are between 0 and 1 and that the measure is symmetric. Following McDaid's et al. definition of $NMI_{LFK}$ (which differs slightly from the formulation of NMI in [1]), for PSTs $X$ and $Y$, a normalized MI can be formulated as (Eq. 7 in [2]):

$$NMI(X,Y) = 1 - \frac{1}{2}\left(\frac{H(X|Y)}{H(X)} + \frac{H(Y|X)}{H(Y)}\right). \tag{S5}$$

This formulation ensures that the measure equals 1 if $X = Y$.

#### An example of NMI comparisons

In this section, we provide an illustrative example of our application of the NMI measure and its computation. Figure S4 shows an example of two PSTs, defined over a set of 14 individuals, $I = \{a, b, c, d, e, f, g, h, i, j, k, l, m, n\}$. In order to compare these PSTs using the NMI measure, the PSTs are represented as a partition of $I$; PST $A$ as partitions of $I$ into the 8 sets A1-A8 in the figure, and PST $B$ as a partition of $I$ into the 9 sets B1-B9. To compute NMI between $A$ and $B$, for each cluster in $A$ we find the cluster most similar to it in $B$ using Eq. S3, and compute the lack of information between the clusters using Eq. S4; the same is done for each cluster in $B$. For example, for cluster A6 in $A$, the conditional entropy is minimized in Eq. S3 by cluster $B6$ in $B$, with $H(A6|B6) = 4.83$. For the clusters that are identical between the PSTs, (A1,B1), (A2,B2), and (A3,B3), we get $H(A1|B1) = H(A2|B2) = H(A3|B3) = 0$, and these pairs of identical clusters minimize the conditional entropy with no contribution to the lack of information between the PSTs.

After all the appropriate conditional entropies are computed, Eqs. S2, S3, and S5 are used to sum and normalize these values. For the two PSTs in Figure S4, we get: $H(A|B) = 22.08$, $H(B|A) = 31.73$, $H(A) = 77.23$, and $H(B) = 89.31$. These are used in Eq. S5 to produce the NMI score between $A$ and $B$, $NMI(A, B) = 0.679$.

We now turn to comparing only a subset of the clusters in the PSTs, the leaves, as presented in the main text. For $A$, the leaves are A4–A8, and for $B$, the leaves are B5–B9 (green clusters in Fig. S4). Whereas for $A$ the union of the leaves is $I$, this is not the case for $B$, since individual $h$ is not assigned to any leaf. In $B$, the cluster closest to the leaves that contains $h$ is B3, and we therefore add $h$ to all leaves descending from B3, which are B6 and B7. Therefore, the partition representation of $B$ which we use to compute NMI for the leaf clusters is $\{a, b\}, \{c, d\}, \{e, f\}, \{g, h, i, j\}, \{h, k, l, m, n\}$. For $A$, we use the sets A4–A8. The NMI value for the comparison between these representations is 0.567. This value is lower than that attained for the comparison of the entire PSTs, because similarity between clusters at the higher hierarchical levels (orange clusters in Fig. S4) was higher, with three identical pairs of clusters, contributing to a higher NMI score.

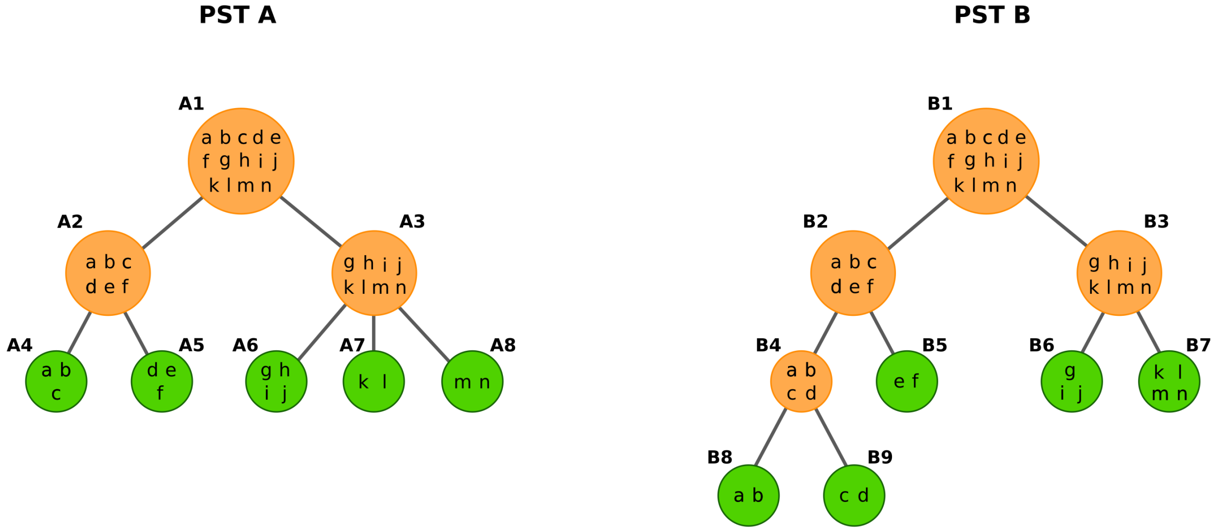


**Figure S5: An example of two PSTs**. Both PSTs are defined over a set $I$ of 14 individuals, marked a–n. Internal clusters are shown in orange, and leaf clusters are shown in green.
